# CVD-associated SNPs with regulatory potential drive pathologic non-coding RNA expression

**DOI:** 10.1101/2023.02.12.528184

**Authors:** Chaonan Zhu, Nina Baumgarten, Meiqian Wu, Yue Wang, Arka Provo Das, Jaskiran Kaur, Fatemeh Behjati Ardakani, Thanh Thuy Duong, Minh Duc Pham, Maria Duda, Stefanie Dimmeler, Ting Yuan, Marcel H. Schulz, Jaya Krishnan

## Abstract

**Background:** Cardiovascular diseases (CVDs) are the leading cause of death worldwide. Genome-wide association studies (GWAS) have identified many single nucleotide polymorphisms (SNPs) appearing in non-coding genomic regions in CVDs. The SNPs may alter gene expression by modifying transcription factor (TF) binding sites and lead to functional consequences in cardiovascular traits or diseases. To understand the underlying molecular mechanisms, it is crucial to identify which variations are involved and how they affect TF binding.

**Methods:** The SNEEP (SNP exploration and analysis using epigenomics data) pipeline was used to identify regulatory SNPs, which alter the binding behavior of TFs and link GWAS SNPs to their potential target genes for six CVDs. The human induced pluripotent stem cells derived cardiomyocytes (hiPSC-CMs), monoculture cardiac organoids (MCOs) and self-organized cardiac organoids (SCOs) were used in the study. Gene expression, cardiomyocyte size and cardiac contractility were assessed.

**Results:** By using our integrative computational pipeline, we identified 1905 regulatory SNPs in CVD GWAS data. These were associated with hundreds of genes, half of them non-coding RNAs (ncRNAs), suggesting novel CVD genes. We experimentally tested 40 CVD-associated non-coding RNAs, among them RP11-98F14.11, RPL23AP92, IGBP1P1, and CTD-2383I20.1, which were upregulated in hiPSC-CMs, MCOs and SCOs under hypoxic conditions. Further experiments showed that IGBP1P1 depletion rescued expression of hypertrophic marker genes, reduced hypoxia-induced cardiomyocyte size and improved hypoxia-reduced cardiac contractility in hiPSC-CMs and MCOs.

**Conclusions:** IGBP1P1 is a novel ncRNA with key regulatory functions in modulating cardiomyocyte size and cardiac function in our disease models. Our data suggest ncRNA IGBP1P1 as a potential therapeutic target to improve cardiac function in CVDs.

## Introduction

Cardiovascular diseases (CVDs) are among the most common causes of death in the world. Finding novel molecular biomarkers is an important research goal, to enable development of novel early detection, treatment and intervention strategies.

Recently, non-coding RNAs (ncRNAs) have been found to play important roles in cellular processes related to many CVDs.^1–3^ The ncRNA HERNA1 (hypoxia-inducible enhancer RNA 1), which is produced by direct hypoxia-inducible factor 1α binding to an hypoxia response element, modulates the cardiac growth, metabolic, and contractile gene program in pressure-overload heart disease.^4^ Similarly, the ncRNA CARMEN is derived from a human super-enhancer (SE) and regulates cardiomyocyte differentiation and homeostasis in human cardiac precursor cells.^5^ Moreover, inhibition of the ncRNA MEG3 (maternally expressed gene 3) decreased cardiac fibrosis and improved diastolic performance by targeting cardiac matrix metalloproteinase-2 (MMP-2).^6^

Different approaches can be used to associate a ncRNA with the pathology of a CVD. Genome-wide methods have been especially successful using different types of assays measuring RNA,^7,8^ genome,^9^ epigenome^10^ variation or image-measured physiological differences^11^ in disease models or from patient data directly.

Genome data in the form of mutations, such as single nucleotide polymorphisms (SNPs), that are associated with CVDs through genome-wide association studies (GWAS), provide an interesting source of information for detection of relevant ncRNAs. In particular, because many mutations found associated with CVDs reside outside of protein-coding genes and their functional role is often unknown.^9^ However, it is difficult to connect SNPs that reside in the non-coding regions of the genome with potential biological functionality.

One promising direction for deciphering the functional role of such non-coding SNPs are variant annotation methods that use transcription factor (TF) binding or epigenetic information such as DNase1-seq, ATAC-seq or histone ChIP-seq data.^12,13^ To predict TF binding, different approaches exist using position weight matrices (PWMs),^14,15^ that are available for the majority of human TFs, or more complex methods such as deep learning based models^16,17^, which are currently more limited due to lack of TF-specific data. Specific statistical methods have been developed to assess whether a SNP has a regulatory effect on TF binding.^18–20^

Alternatively, SNPs can be categorized as *functionally important* by a computational model that assesses whether changes in the DNA sequence will affect gene or epigenome activity more generally.^21–23^ In other words, all these methods assess whether a SNP is likely to have a regulatory effect and may allow to predict the tissue and cell-type relevance of such an effect.^24–26^

While prioritization of regulatory SNPs (rSNPs) with approaches mentioned above is important and an area of active research, another problem is to associate rSNPs with their potential target genes. Several approaches for linking regions to target genes exist using diverse data types,^27,28^ such as the Activity-by-Contact model^29^ or STITCHIT.^30^ For example, the EpiRegio database^31^ contains 2.4 million regulatory elements (REMs) that were linked to human target genes using STITCHIT utilizing paired DNase1-seq and RNA-seq data of several cell types.

Here, we present a characterization of ncRNAs that can be linked to genetic mutations associated with the CVDs, including Aortic stenosis, Coronary artery disease, Cardiomyopathy, Cardiac arrhythmia, Myocardial infarction or Myocardial ischemia. By using an algorithm to detect rSNPs as part of the SNEEP pipeline, hundreds of cardiovascular associated ncRNAs have been identified that harbor rSNPs in their gene-regulatory elements. To study the functions of some interesting ncRNAs we used two models: the 2D human induced pluripotent stem cells (hiPSCs) derived cardiomyocytes (hiPSC-CMs) model and 3D human cardiac organoids, which display a similar microenvironment and contractile function to the human heart. Through assessing the cardiomyocyte size and contractile function response to pathophysiologic stress, our data demonstrated that ncRNA immunoglobulin (CD79A) binding protein 1 pseudogene 1 (IGBP1P1) drives cardiac hypertrophy and contractile dysfunction.

## Methods

### Collection of GWAS SNPs of cardiovascular diseases

We have collected the significant SNPs from the NHGRI-EBI GWAS catalog^32^ for the following search terms including all the available child traits: Coronary artery disease (EFO_0000378), Aortic stenosis (EFO_0000266), Cardiac arrhythmia (EFO_0004269), Cardiomyopathy (EFO_0000407), Myocardial infarction (EFO0000612) and Myocardial ischemia (EFO0005672). All GWAS were downloaded on 10/26/2020 (see also Table S5).

For each set of GWAS SNPs, we have obtained correlated SNPs that are in linkage disequilibrium (LD) with any of the original SNPs. We used the LDProxy Tool^33^ and extracted the proxy SNPs via their API functionality. Proxy SNPs with an R^2^ >= 0.75 and within a window of +/- 500 000 bp centered around the original SNP were added to the GWAS SNPs. The combined set of proxy and lead SNPs was used as input set to SNEEP.

### Detection of regulatory SNPs

To detect regulatory SNPs, we applied the SNEEP pipeline (https://github.com/SchulzLab/SNEEP) separately for each of the 6 cardiac GWAS. SNEEP (SNP exploration and analysis using epigenomics data) is a computational pipeline which identifies rSNPs along with the affected TFs and further links the rSNPs to putative target genes.

To compute whether or not a SNP alters the binding behavior of a TF, a differential binding score is determined, which is the log-odds ratio between the binding affinity of the wildtype sequence (containing the wild type allele) and the mutated sequence (containing the alternative allele).^20^ For the log-odds ratio a differential binding p-value is computed. The approximation of the p-value depends on the characteristics like length, CG content etc. of the used TF PWM-motifs. Therefore, one needs to estimate a scale value per motif using the script *estimateScalePerMotif*.*sh* from the SNEEP pipeline. To estimate the scale parameter values, we used 200 000 sampled SNPs from the dbSNP database^34^, and removed flanking bases of the PWMs with an entropy higher than 1.9, resulting in the following command: bash estimateScalePerMotif.sh 200000 <pathToMotifs> <outputDir> <motifNames> 1.9

To run SNEEP, 632 human TF PWM-motifs in transfac format were gathered from the JASPAR database (version 2020)^35^ and over 2.4 million regulatory elements linked to their putative target genes were downloaded from the EpiRegio database^31^ (https://doi.org/10.5281/zenodo.3758494,file: REMAnnotationModelScore_1.csv.gz).

Next, we applied the main SNEEP pipeline per GWAS with a differential binding p-value cutoff of 0.001:

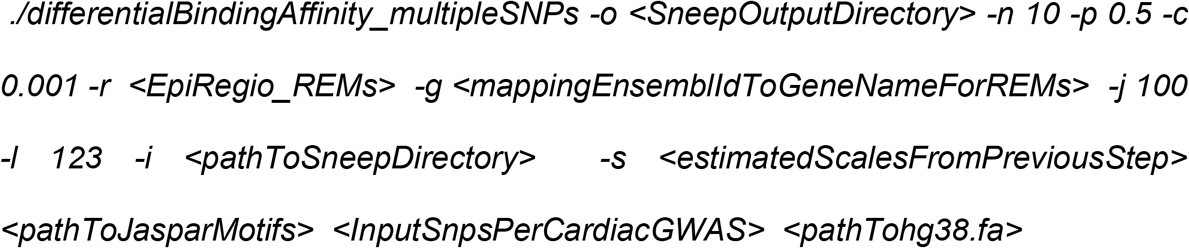

The SNEEP result is provided in Table S6.

### Identification of disease associated genes using rSNPs

As part of the analysis of SNEEP, all rSNPs that overlap regulatory elements from Epiregio constitute a candidate disease gene. From the SNEEP output file protein-coding and non-coding genes were extracted that have overlapping rSNPs in their regulatory elements. Non-coding genes were understood to be all genes not labeled with the biotype ‘protein coding’ or ‘TEC’ (primary assembly annotation, version 39, downloaded from GENCODE).^36^

To label which protein-coding genes are already associated with the studied diseases (Figure 2B, black dots), we used the *disease2gene* functionality of the R package of DisGeNET:

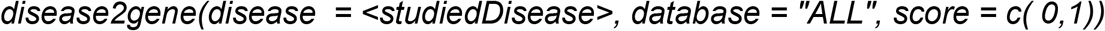

The *studiedDisease* parameter needs to be provided as UMLS CUI identifiers. We used C1956346 (CAD), C0003811 (Cardiac arrhythmia), C1449563 (Cardiomyopathy, Familial Idiopathic), C0151744 (Myocardial ischemia), C0027051 (Myocardial infarction) and C0340375 (Subaortic stenosis) as *studiedDisease* (see also Table S2).

### Identification of co-expressed genes for non-coding genes and disease enrichment

We conducted a co-expression analysis using the gene expression profiles of 9,662 GTEx RNA-seq samples.^37^ We compared the expression of protein-coding genes with the non-coding genes using the Spearman correlation coefficient as the similarity metric (Table S7). This allowed us to obtain a ranked list of protein-coding genes most similar to the expression of a selected non-coding gene in the GTEx data.

For each non-coding gene the top 10 co-expressed protein coding genes were extracted, varying this number did not change the further results. The joint set of all protein-coding genes that are co-expressed to any of the non-coding genes found for the same disease via the GWAS analysis, were considered to perform a disease enrichment analysis. The analysis was done separately for the resulting co-expressed protein-coding gene sets derived from Cardiac arrhythmia, CAD and Cardiomyopathy using the function *disease_enrichment* from the R package of DisGeNET.^38^

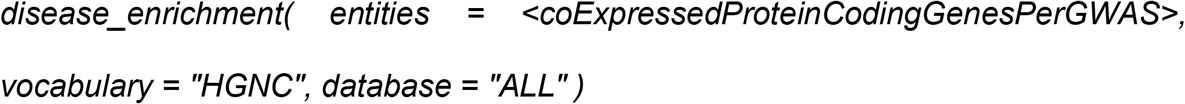

From the resulting list of enriched diseases (Table S3), as part of the enrichment computation, cardiac phenotypes were selected and visualized as a dot plot using ggplot2 (Figure 3C). For the GWAS Aortic stenosis, Myocardial infarction and Myocardial ischemia we did not apply the disease enrichment analysis because of the low number of associated non-coding genes (less than 30).

### Preparation and maintenance of human iPSC-CMs

Human induced pluripotent stem cells (hiPSCs) were purchased from Cellular Dynamics International (CMC-100-010-001) and cultured according to the manufacture’s protocol. The human iPSC-derived cardiomyocytes (hiPSC-CMs) were reprogrammed using the STEMdiff^™^ Cardiomyocyte Differentiation Kit (STEMCELL Technologies) as recommended by the manufacturer. Briefly, human iPSCs were plated at cell density of 3.5×10^5^ cells/well on Matrigel coated 12-well-plates using mTeSR^™^ medium supplemented with 5µM ROCK inhibitor (Y-27632, STEMCELL Technologies) for 24 h. After 1 day (−1), the medium was replaced with fresh TeSR™ medium. To induce cardiac differentiation, the TeSR™ medium was replaced with Medium A (STEMdiff™ Cardiomyocyte Differentiation Basal Medium containing Supplement A) at day 0, Medium B (STEMdiff™ Cardiomyocyte Differentiation Basal Medium containing Supplement B) at day 2, Medium C (STEMdiff™ Cardiomyocyte Differentiation Basal Medium containing Supplement C) at day 4 and day 6. On day 8, medium was switched to STEMdiff™ Cardiomyocyte Maintenance Medium with full medium changes every 2 days, to promote further differentiation into mature cardiomyocyte cells. All experiments were performed in the hiPSC-CMs at day 40. Hypoxic condition was achieved by using the Hypoxia chamber and the hiPSC-CMs were cultured at either 3% or 1% O_2_ for 2 days.

### Monoculture cardiac organoid formation Technique

Monoculture cardiac organoids (MCOs) were created by hiPSC-CMs. Aggrewell^™^ 800 microwell culture plates were used to create the MCOs in STEMdiff™ Cardiomyocyte Support Medium (STEMCELL Technologies). At day 18, hiPSC-CMs were distributed into Aggrewell^™^ 800 microwell culture plates at a density of 900,000 hiPSC-CMs/well. After 2 days of culture, medium was switched to STEMdiff™ Cardiomyocyte Maintenance Medium for long term culture. Hypoxic condition was achieved by using the Hypoxia chamber and the MCOs were cultured at either 3% or 1% O_2_ for 3 days.

### Self-organized Cardiac Organoids formation Technique

Human iPSCs were plated at cell density of 1.5×10^5^ cells/well on Aggrewell^™^ 800 microwell culture plates to form embryoid bodies (EBs). At day 18 and day 20, 50 nM VEGF and 25 nM FGF were added into the Maintenance Medium. On day 22, medium was switched to Maintenance Medium with EGM-2 with full medium changes every 2 days. All experiments were performed in the self-organized cardiac organoids (SCOs) at day 40. Hypoxic condition was achieved by using the Hypoxia chamber and the SCOs were cultured at either 3% or 1% O_2_ for 3 days.

### RNA isolation, reverse transcription and qRT-PCR

Samples were harvested in QIAzol Lysis Reagent (QIAGEN), and total mRNAs were isolated with RNeasy Kit (QIAGEN) according to the manufacturer’s protocol. 200ng total RNAs were reverse transcript into cDNA using QuantiTect Reverse Transcription Kit (QIAGEN) according to the manufacturer’s protocol. The Applied Biosystems StepOnePlus Real-Time PCR system (Applied Biosystems, CA, USA) with Fast SYBR Green Master Mix (Thermo Fisher Scientific) were used for analysis. Gene expression levels were normalized against the housekeeping gene *HPRT1*. The qRT-PCR primers are listed in Table S9.

### Antisense LNA GapmeRs

Antisense LNA GapmeRs were purchased from Qiagen. Four different antisense LNA GapmeRs were designed for each target (Table S10). The GapmeR negative control was used as the control.

### Contractility measurement

Every single MCO was transferred into 96-well-plate and treated with either GapmeR control, GapmeR RP11-98F14.11, GapmeR RPL23AP92 or GapmeR IGBP1P1. Hypoxic condition was achieved by using the Hypoxia chamber and cardiac organoids were cultured at 3% O_2_ for 3 days, then the contractility will be analyzed by IonOptix system. Units/pixels were determined by calibrating the system with a micrometer.

### Calcium Transient measurements

The MCOs were cultured in the 96-welll-plate. 2µM Cal-520 AM (AAT bioquest) with 0.04% Pluronic® F-127 (AAT bioquest) working solution was added into the plate, and then the plate was incubated in the incubator at 37 °C for 90 min. After washing the MCOs with PBS for 3 times, and with medium for 2 times, the MCOs will be transferred to the 384 well U-bottom plate and incubated at 37 °C for 1 h. The fluorescence will be measured by the fluorescence plate reader.

### Immunofluorescence staining

Immunofluorescence staining was performed as described previously. After fixation with 4% paraformaldehyde (PFA)/PBS, the hiPSC-CMs or MCOs were permeabilized and incubated overnight at 4°C with primary antibodies against sarcomeric α-actinin (Sigma Aldrich), diluted in 2% (v/v) HS/PBS. After 3 washes with PBS for 5 min, cells were incubated with 4’,6-diamidino-2-phenylindole (DAPI Thermo Fisher Scientific) and AlexaFluor 555 anti-mouse (Thermo Fisher Scientific) secondary antibody for 1 h at room temperature. Dishes were mounted onto glass slides (Fisher Scientific) with a drop of ProLong™ Gold Antifade (Thermo Fisher Scientific). Fluorescent images were acquired with the SP5 confocal microscopy (Leica) using a 40x magnification. Cell size was quantified blindly using the software Image J.

### Statistical Analysis

Data are represented as mean and error bars indicate the standard error of the mean (SEM). Two-tailed unpaired Student’s *t*-tests (Excel) or one-way ANOVA analyses followed by either a Dunnett’s multiple comparison post-test (multiple comparisons to a single control) or Bonferroni correction (multiple comparisons between different groups) were used as indicated in the respective figure legends. ns= not significant; \**P*< 0.05; \*\**P*< 0.01; %*P*< 0.05.

## Results

### Identification of cardiovascular disease associated genes using regulatory SNPs and enhancer-gene linkage

Based on the previously postulated idea to identify rSNPs that have a regulatory effect, we have used the SNEEP pipeline to combine three sources of information for finding genes related to a CVD: 1) SNPs found significantly associated in GWAS with a particular CVD,^32^ 2) prediction which of these SNPs are potentially regulatory using human TF PWM-motifs,^20,35^ and 3) enhancer-gene catalogue to map SNPs to putative target genes (Figure 1).^31^ In short, SNEEP uses PWM descriptions of TF binding sites to assess whether a SNP would affect the binding of a known TF. Such an rSNP may lead to the loss of a TF binding site or the creation of a new binding site affecting the expression of one or several target genes related to the phenotype.

**Figure 1.**
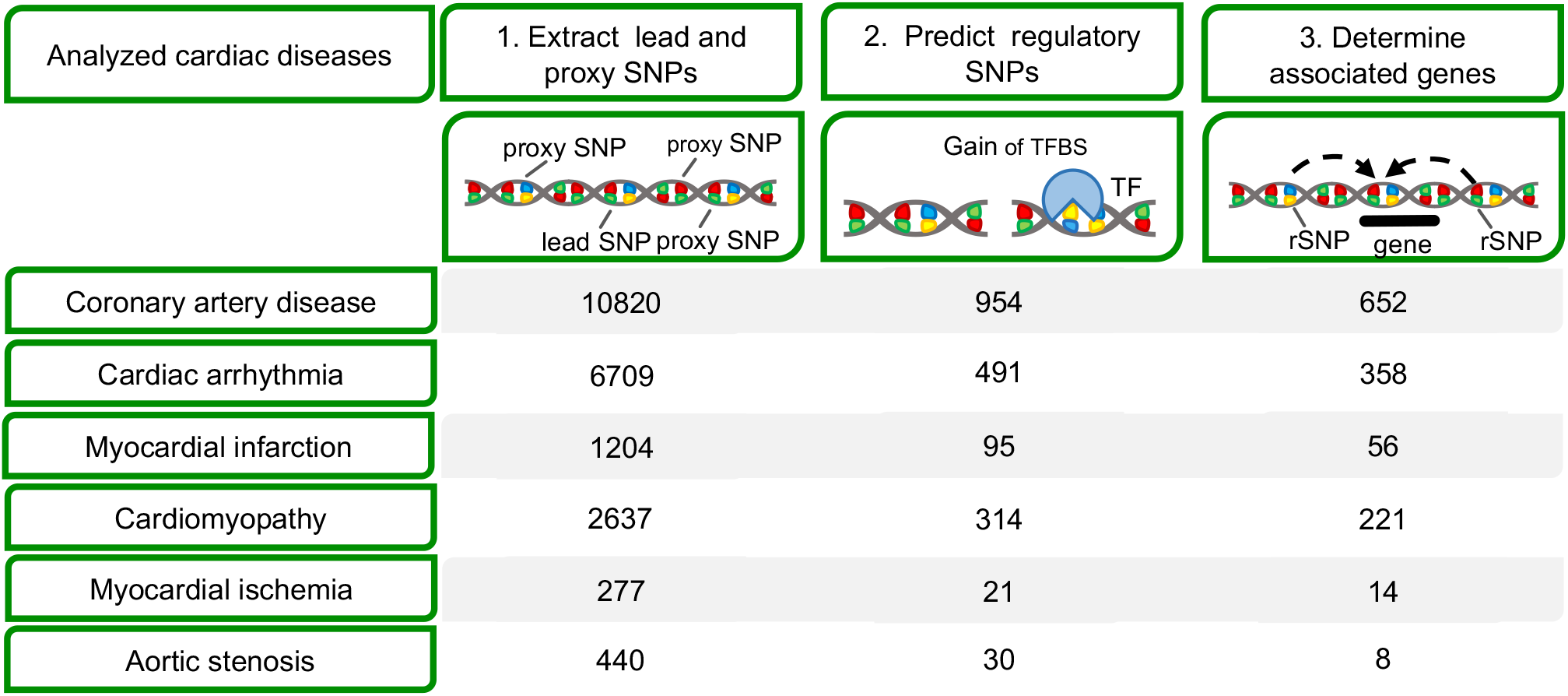
Overview of the bioinformatic pipeline to identify non-coding genes associated with cardiovascular diseases. Step 1: SNPs for 6 different cardiac diseases were collected from the NHGRI-EBI GWAS catalog. Step 2: We used existing transcription factor binding models to filter regulatory SNPs (rSNPs), that are predicted to have an impact on transcription factor binding sites (TFBS). Step 3: rSNPs are linked to putative target genes using enhancer-gene links. For each step the number of SNPs or genes for each disease is given per row.

In the first step, we have retrieved significant lead SNPs and correlated SNPs in LD (R^2^ >= 0.75) from GWAS for known cardiovascular diseases: 10,820 for Coronary artery disease (CAD), 440 for Aortic stenosis, 2,637 for Cardiomyopathy, 6,709 for Cardiac arrhythmia, 1,204 for Myocardial infarction and 277 for Myocardial ischemia, which were then subjected to SNEEP analysis (Figure 1). First, we filtered for those SNPs that may have a regulatory function (rSNPs) and affect TF binding. Then, we utilized the EpiRegio database, containing 2.4 million human regulatory elements, to link rSNPs to possible target genes for the respective indications.

These analyses led to the identification of protein-coding and non-coding genes that could be directly linked to a specific indication (Figure 2A). Notably, there were often similarly many non-coding and protein-coding genes associated. We used the DisGeNET database^38^, a large collection of known disease associated genes, for a positive control experiment. We conducted a disease enrichment analysis that assessed whether the newly identified protein-coding genes using our approach are enriched among previously associated disease genes. We found that for all tested indications we were able to get the corresponding disease phenotype as significantly enriched (Fisher’s exact test, FDR <= 0.05, Figure S1, Table S1). For Myocardial ischemia and Aortic stenosis, we had less than 30 genes available and therefore enrichment analysis was omitted as it is statistically underpowered. The enrichment analysis results support that our approach, although limited to using rSNPs, is able to find many of the previously associated disease genes.

**Figure 2.**
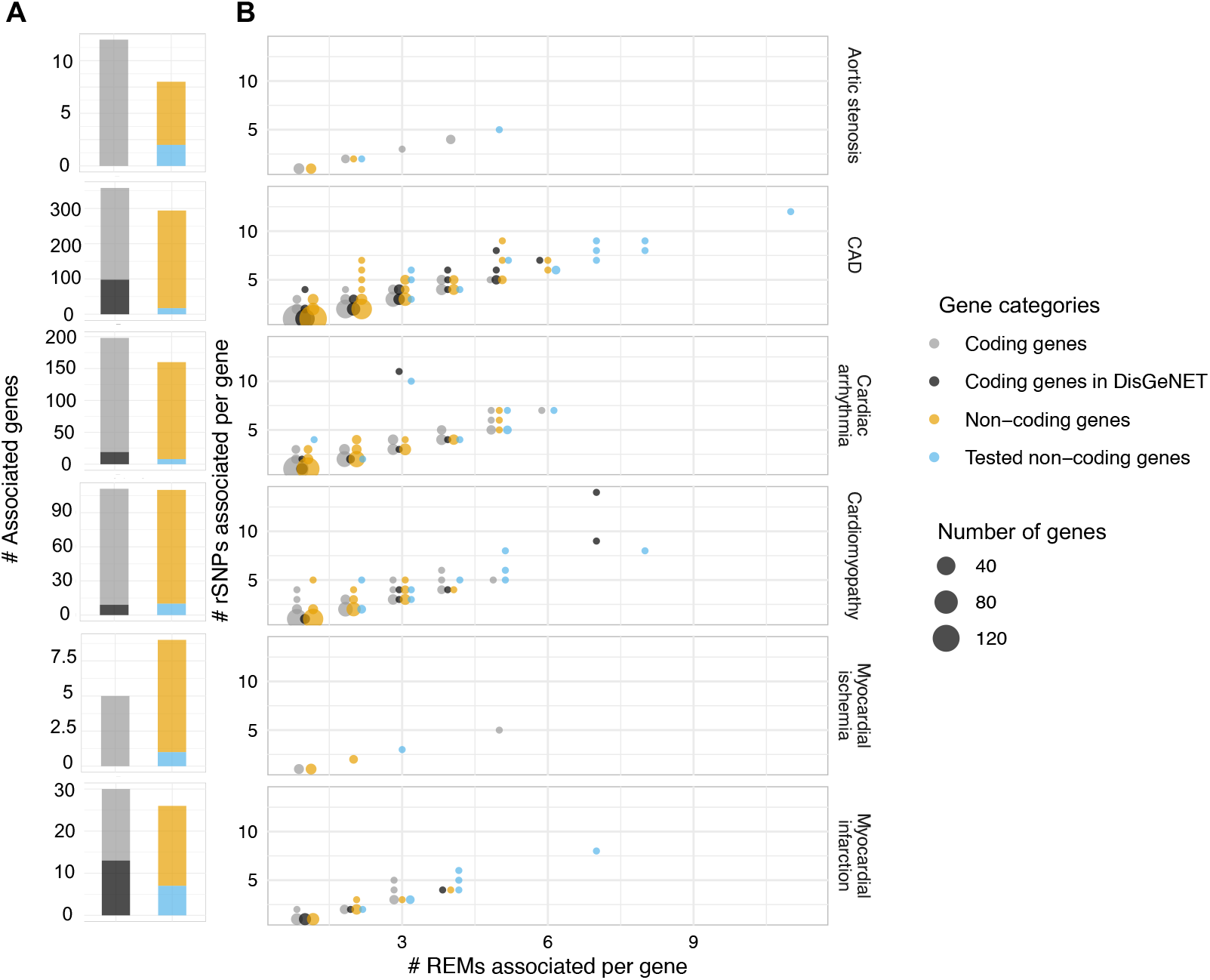
Analysis of protein and non-coding genes associated with rSNPs. (A) For each GWAS (row) a stacked bar plot shows the number of associated genes per category. (B) Dot plot visualizing per GWAS how many regulatory elements (REMs) and rSNPs are associated with a gene. Genes are separated into protein-coding, protein-coding associated with the disease according to DisGeNET, non-coding, and experimentally studied non-coding genes in this work (circle colour). The x-axis shows the number of REMs for a gene overlapping with at least one rSNP and the y-axis the number of rSNPs for all REMs associated with a gene. The size of the dot correlates with the number of genes having the same x- and y-coordinate values.

We systematically compared the properties of the associated genes, looking at the number of rSNPs that could be linked to each gene and the number of regulatory elements of each gene with at least one rSNP (Figure 2B, Table S2). Protein- and non-coding genes showed similar characteristics. Notably, identified genes had on average more rSNPs than disease genes listed in DisGeNET. This underlines a unique feature of our approach, highlighting genes that have many non-coding rSNPs associated with the indication. Of particular note was the large number of non-coding genes we identified. We surveyed the literature and existing databases that list known RNA biomarkers to identify additional evidence for genes that we found in our analyses (Table S2). For example, we checked the Heart Failure database for known RNA biomarkers (HFBD),^39^ but none of the 49 ncRNAs listed there overlapped ours.

In speculating that it would be possible to find further evidence using existing OMICs data - we used a guilt-by association strategy and collected for each non-coding gene the top 10 most correlated protein-coding genes, according to a large RNA expression dataset from the GTEx resource^37^ (Figure 3A). We then gathered all co-expressed protein-coding genes with respect to the non-coding genes we had found for each disease (Figure 3B). This collection thus signifies all protein-coding genes that are highly correlated to the non-coding genes we found. We conducted the DisGeNET enrichment analysis for all coexpressed protein-coding genes (Figure 3C, Table S3). We observed a strong enrichment for the expected diseases in each tested study in which we had more than 30 non-coding genes. Thus, we were positive that a majority of the non-coding genes could play an important role in the underlying disease.

**Figure 3.**
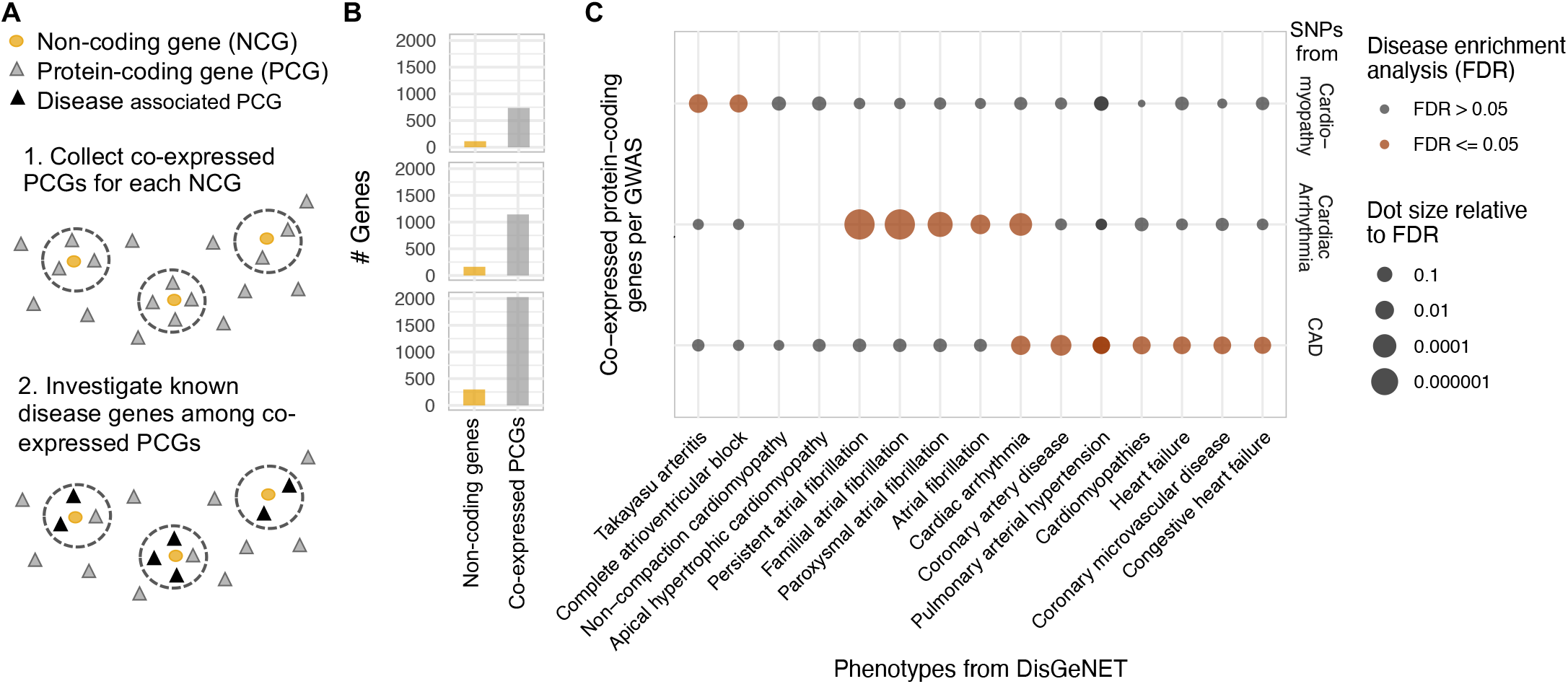
Disease enrichment analysis based on the protein coding genes co-expressed with GWAS-associated non-coding genes. (A) We sought to identify co-expressed protein-coding genes (PCGs, triangles) for each non-coding gene (NCG, circle) using a large dataset of RNA-seq samples from GTEx^34^. The top correlated protein-coding genes are tested for an enrichment of previous indications with a cardiovascular disease. (B) Barplot visualizing the number of non-coding and co-expressed protein-coding genes per GWAS (row). (C) Dot plot showing enriched cardiovascular phenotypes from DisGeNET (x-axis) for the top 10 co-expressed protein-coding genes of the associated non-coding genes separated per GWAS (y-axis). The dot coloring represents whether a phenotype is significantly enriched (FDR<= 0.05) and the dot size is relative to the FDR.

### Identification and characterization of ncRNAs in hiPSC-CMs

Our previous analyses suggested that many of the ncRNAs that we associated with CVDs could have important cardiovascular functions. Thus, we selected in total 40 ncRNA genes and prioritized ncRNA genes not listed in DisGeNET and with many rSNPs (Figure 2B, Table S4 and S8). To determine the roles of the 40 ncRNAs in human cardiomyocytes, the hiPSC derived CMs, monoculture cardiac organoids (MCOs) and self-organized cardiac organoids (SCOs) were used in the study (Figure 4A). The hiPSC-CMs, MCOs and SCOs were incubated in normoxia and hypoxia (1% O_2_ or 3% O_2_, respectively), and the expression levels of these 40 ncRNAs were quantified by qRT-PCR (Figure 4B). As shown in Figure 4B, we have selected the ncRNAs RP11-98F14.11, RPL23AP92, IGBP1P1, and CTD-2383I20.1 for further experiments, as they are consistently upregulated in hypoxia in hiPSC-CMs, MCOs and SCOs.

**Figure 4.**
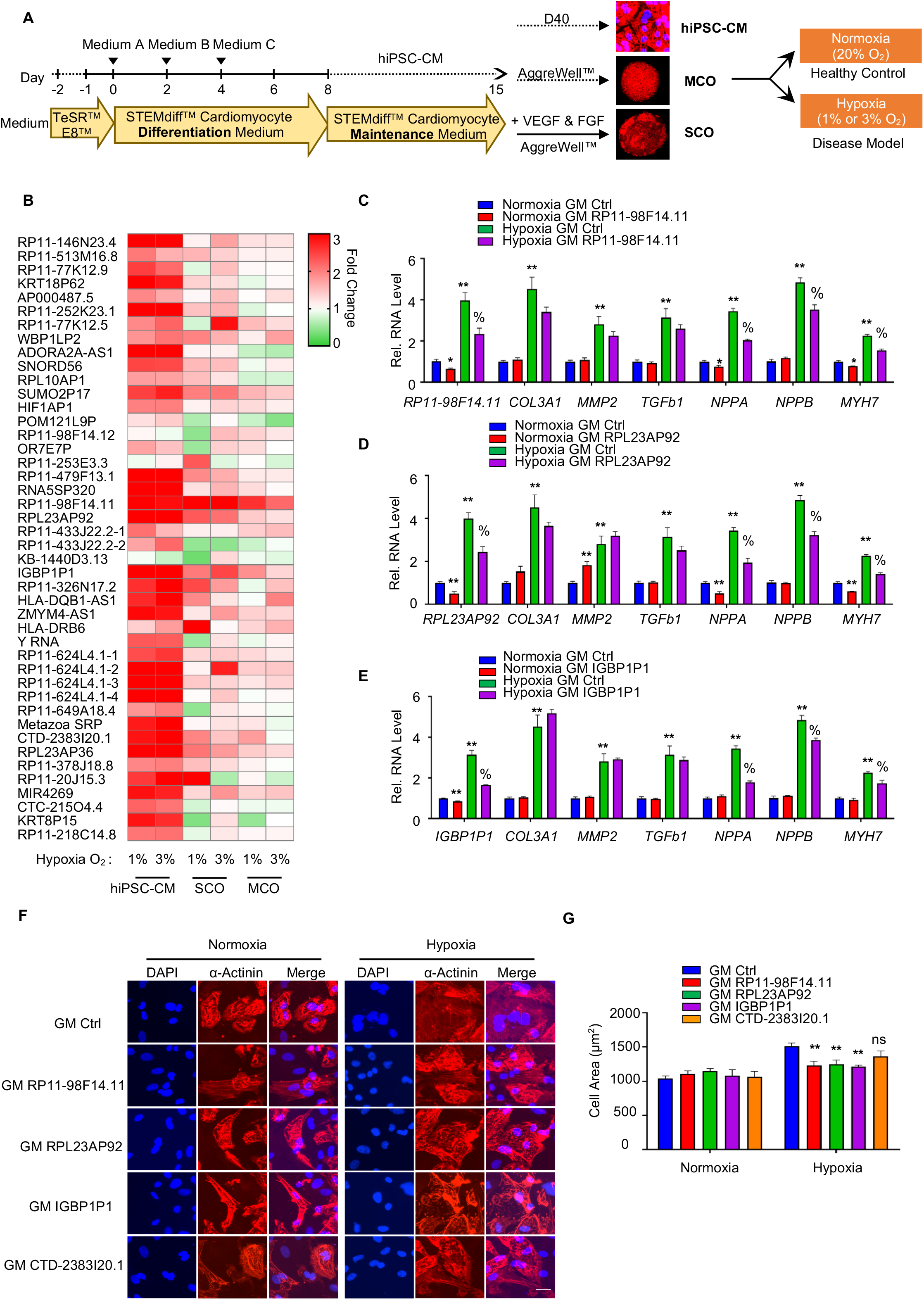
Identification and characterization of ncRNAs in hiPSC-CMs. (A) Schematic protocol for differentiation of hiPS cells into cardiomyocytes, monoculture cardiac organoids (MCOs) and self-organized cardiac organoids (SCOs) *in vitro*. (B) Heat map shows the relative mRNA expression levels of the 40 ncRNAs in the 40-day-old hiPSC-CMs, or 40-day-old human cardiac organoids (including MCOs and SCOs) treated with either 1% O_2_ or 3% O_2_ for 2 or 3 days, respectively. Data are normalized to either normoxia hiPSC-CMs, or normoxia MCOs/SCOs. The red and green colors indicate high and low expression values, respectively. Means of n=3 biological replicates per group. (C) Relative mRNA expression of *RP11-98F14*.*11, COL3A1, MMP2, TGFb1, NPPA, NPPB* and *MYH7* in hiPSC-CMs transduced with either GapmeR RP11-98F14.11 (GM RP11-98F14.11) or GapmeR negative control (GM Ctrl) in both normoxia and hypoxia. Data are represented as Mean±SEM; n=3; *, %*P*< 0.05. (D) Relative mRNA expression of *RPL23AP9, COL3A1, MMP2, TGFb1, NPPA, NPPB* and *MYH7* in hiPSC-CMs transduced with either GapmeR RPL23AP9 (GM RPL23AP9) or GM Ctrl in both normoxia and hypoxia. Data are represented as Mean±SEM; n=3; *, %*P*< 0.05. (E) Relative mRNA expression of *IGBP1P1, COL3A1, MMP2, TGFb1, NPPA, NPPB* and *MYH7* in hiPSC-CMs transduced with either GapmeR IGBP1P1 (GM IGBP1P1) or GM Ctrl in both normoxia and hypoxia. Data are represented as Mean±SEM; n=3; *, %*P*< 0.05. (F) Representative images of α-actinin (red) and DAPI (blue) in 40-day-old hiPSC-CMs after GM RP11-98F14.11, GM RPL23AP9, GM IGBP1P1, GM CTD-2383I20.1, or the GM Ctrl under normoxia or hypoxia for 2 days. Scale bar is 25 µm. (G) Cell size of the hiPSC-CMs was assessed by the Image J. 50 cells were analyzed for each condition. Data are represented as Mean±SEM; n=3; *, %*P*< 0.05. hiPSC-CMs, human induced pluripotent stem cell-derived cardiomyocytes; MCOs, monoculture cardiac organoids; SCOs: Self-organized cardiac orgnoids.

In order to investigate the function of RP11-98F14.11, RPL23AP92, IGBP1P1, and CTD-2383I20.1 in hiPSC-CMs, we performed loss-of-function through GapmeR-mediated knockdown of the respective target ncRNAs. GapmeRs are chimeric anti-sense oligonucleotides that contain a central block of deoxynucleotide monomers to induce RNaseH cleavage.^40^ Four GapmeRs were designed and synthesized for each ncRNA target (Table S10), and the knockdown efficiency was quantified by qRT-PCR in hiPSC-CMs (Figure S2A-S2D). We selected GapmeRs RP11-98F14.11 #2 (GM RP11-98F14.11), GapmeR RPL23AP92 #1 (GM RPL23AP92), GapmeR IGBP1P1 #1 (GM IGBP1P1), and GapmeR CTD-2383I20.1 #1 (GM CTD-2383I20.1) for further experiments. As shown in Figure 4C, the expression levels of pathologic hypertrophy markers, including atrial natriuretic peptide A (*NPPA*), *NPPB* and beta-myosin heavy chain 7 (*MYH7*), were reduced after GapmeR RP11-98F14.11 (GM RP11-98F14.11) treatment in hypoxia compared to the GapmeR Control (GM Ctrl) treatment. We also profiled expression of key fibrotic marker genes including Collagen type III alpha 1 chain (*COL3A1*), matrix metalloproteinase-2 (*MMP2*) and Transforming growth factor β1 (*TGFb1*). However, GM RP11-98F14.11 treatment did not alter hypoxia induced fibrotic marker gene expression in hiPSC-CMs compared to the GM Ctrl. Similarly, Knockdown of RPL23AP92 and IGBP1P1 reduced hypoxia-induced pathologic hypertrophy marker expression, but not of hypoxia-induced fibrotic marker genes (Figure 4D and 4E). CTD-2383I20.1 did not alter either hypoxia-induced pathologic hypertrophy markers or hypoxia-induced fibrotic marker gene expression compared to GM Ctrl (Figure S3).

To better define the function of RP11-98F14.11, RPL23AP92, IGBP1P1, and CTD-2383I20.1 in human cardiomyocytes, we analyzed cardiomyocyte cell size in a loss-of-function setting. In order to directly visualize the cells, cardiomyocytes were stained for α-actinin and DAPI and imaged by confocal microscopy (Figure 4F). As shown in Figure 4G, hypoxia led to increased cell size, which was rescued upon RP11-98F14.11, RPL23AP92 and IGBP1P1 inhibition, while CTD-2383I20.1 did not affect the cell size. Together, this suggests that RP11-98F14.11, RPL23AP92, IGBP1P1 play key roles in regulation of cell size to determine hypertrophic response in human cardiomyocytes, but not in pathology of fibrosis in 2D monolayer cultures.

### IGBP1P1 drives pathologic hypertrophy and contractile dysfunction in human cardiac tissue mimetics

We determined the function of RP11-98F14.11, RPL23AP92, IGBP1P1 in human iPSC-derived cardiac organoids, serving as a model for native heart tissue. On day 18 of differentiation, the monolayer was dissociated into single cells and seeded on Aggrewell^™^ 800 microwell culture plates in order to induce the formation of cardiac organoids (Figure 4A). Cardiac mimetics were then cultured under control normoxic conditions, or in hypoxia (3% O_2_) to mimic myocardial hypoxia *in vitro*. As shown in Figure 5A, hypoxia-induced pathologic hypertrophy markers and fibrotic marker genes were decreased by inhibition of RP11-98F14.11. Similarly, IGBP1P1 depletion also reduced the expression levels of pathologic hypertrophy markers and fibrotic marker genes in hypoxia. In addition, the cell size of cardiomyocytes in MCOs was also measured in both normoxia and hypoxia (Figure 5B). As shown in Figures 5C and 5D, inhibition of IGBP1P1 reduced the cell size in hypoxia compared to the GM Ctrl. However, RPL23AP92 deletion did not affect pathologic hypertrophy markers and fibrotic marker genes (data not shown).

**Figure 5.**
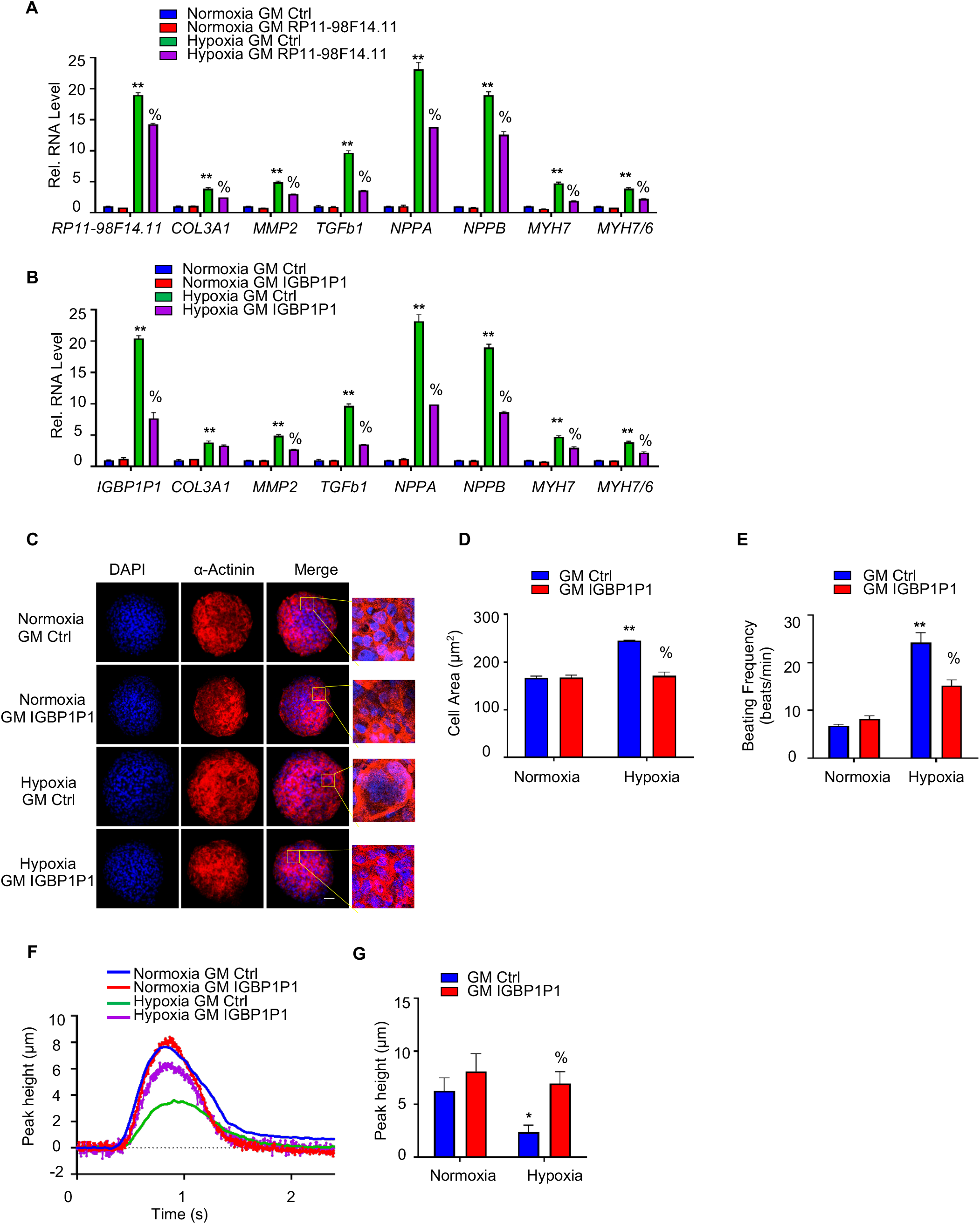
IGBP1P1 is vital in cell size and contractility in human cardiac tissue mimetics. (A) Relative mRNA expression of *RP11-98F14*.*11, COL3A1, MMP2, TGFb1, NPPA, NPPB, MYH7* and ratio *MYH7/6* in MCOs transduced with either GM RP11-98F14.11 or GM Ctrl in both normoxia and hypoxia. Data are represented as Mean±SEM; n=3; *, %*P*< 0.05. (B) Relative mRNA expression of *IGBP1P1, COL3A1, MMP2, TGFb1, NPPA, NPPB, MYH7* and ratio *MYH7/6* in MCOs transduced with either GM IGBP1P1 or GM Ctrl in both normoxia and hypoxia. Data are represented as Mean±SEM; n=3; *, %*P*< 0.05. (C) Representative images of α-actinin (red) and DAPI (blue) in 40-day-old human MCOs after GM IGBP1P1 or the GM Ctrl in normoxia or hypoxia for 3 days. Scale bar is 50 µm. (D) Cardiomyocyte cell size in the MCOs was assessed by the Image J. 50 cells were analyzed for each condition. Data are represented as Mean±SEM; n=3; *, %*P*< 0.05. (E) Human MCOs were treated with GM IGBP1P1 or GM Ctrl in both normoxia and hypoxia for 3 days, and then contraction was determined by counting beats per minute. Data are represented as Mean±SEM; *, %*P*< 0.05. (F) Representative traces of contractile MCOs. (G) The contractility assays were performed by determining the amplitude peak of contracting MCOs. Data are represented as Mean±SEM; *, %*P*< 0.05. All by one-way ANOVA analyses followed by a Dunnett’s multiple comparison post-test. MCOs, monoculture cardiac organoids.

Furthermore, cardiac contractility was also measured by IonOptix and through calcium flux analysis. Both assays revealed that the Hypoxia-induced beating frequency was reduced by IGBP1P1 inhibition (Figure 5E), while hypoxia-reduced contractile amplitude was rescued by IGBP1P1 inhibition compared to GM Ctrl (Figure 5F and 5G, Figure S4A and S4B). However, there is no difference of beating frequency and contractile amplitude between GM RP11-98F14.11 and GM Ctrl (Figure S4C and S4D). Together, IGBP1P1 regulates both cell size and cardiomyocyte contractility in 3D human cardiac tissue mimetics.

## Discussion and Conclusion

In this study we have used known position-weight matrix models of TF binding to predict whether SNPs have a regulatory effect. These models do not consider dependency between positions. Although alternative models exist, such as SLIM,^41^ the absolute number of available TFs is lower than what we used here from the JASPAR database. Exploring more complex models may increase the number of rSNPs that can be detected and thus may reveal additional genes of interest. Instead of predicting rSNPs after the GWAS, Arloth et al. have first determined rSNPs to filter.^26^ They then recomputed the association significance for a smaller subset of genome-wide rSNPs, which may further boost the ability to detect disease genes.

Pseudogenes are defined as regions of the genome that contain defective copies of genes and are often considered as nonfunctional ncRNA. However, recent studies have shown that pseudogenes may play important roles in CVDs.^42^ Elevated low-density lipoprotein cholesterol (LDL-C) level is a main risk factor for CVDs, while knockdown of zinc finger protein 542 pseudogene (ZNF542P) increases the LDL-C level response to simvastatin in a human hepatoma cell line.^43^ The mRNA levels of octamer-binding transcription factor 4 (Oct 4) pseudogene Oct-4-psG1 and Oct-4-psG5 are significantly down-regulated in pulmonary arterial smooth muscle cells (PASMC) in patients with idiopathic pulmonary arterial hypertension (IPAH), indicating that Oct-4-psG1 and Oct-4-psG5 are involved in IPAH.^44^ Moreover, expression level of NMRA-like protein NMRAL1 pseudogene (NMRAL2P) is significantly decreased in the right ventricle in heart failure, suggesting that NMRAL2P is involved in heart failure.^45^ Together, these studies indicate that pseudogenes are closely related to CVDs. IGBP1P1 is a pseudogene of IGBP1, a phosphoprotein associated with the B cell receptor complex and leads to multiple signal transduction pathways.^46^ IGBP1 is a novel biomarker in lupus nephritis (LN) patients, its expression level is increased in the plasma and urine of patients with LN compared with systemic lupus erythematosus (SLE) patients without nephritis and healthy controls.^47,48^ Recently it also showed that IGBP1 is upregulated in esophageal squamous cell carcinoma (ESCC), and its expression is significantly associated with ESCC patient survival.^49^ IGBP1 is also expressed in the heart, but its function has not been studied in the heart yet. In our study, we have identified that IGBP1P1 is upregulated in the human disease model *in vitro*, and its depletion could reduce hypoxia-induced cell size and improve cardiac contractility in human cardiac tissue mimetics. It may modulate cardiomyocyte size and cardiac contractility through regulating the expression level of IGBP1 at both transcriptional and translational levels, but the mechanism is not known yet.

Many studies have demonstrated that ncRNAs play important roles in the development of CVDs.^50^ However, the study of human-specific ncRNAs has been limiting and challenging. The majority of the human lncRNAs are poorly conserved in mouse, conventional mouse models are not a suitable tool to study their function *in vivo* regulation and function. Here, we utilized a human 3D cardiac organoid model, in order to best recapitulate the biological and molecular properties of native heart tissue thus enabling us to study ncRNA function in a physiologically relevant context – lending greater credence to the validity of our study and its findings. The 3D human cardiac organoids are composed of different cell types including cardiomyocytes, endothelial cells, and fibroblasts, which are able to self-organize into complex organ-like structures and have a similar microenvironment to the human heart. In addition, the human cardiac organoids can be cultured for longer term *in vitro*, and also display molecular, metabolic and contractile characteristics of adult native myocardium and respond to pathophysiologic stressors (Figure 5F and 5G, Figure S4A and S4B). The organoid model is a useful biological tool to study the biological functions of ncRNAs in our study. In the future, human cardiac organoids are also the good model for the research in disease modeling, developmental biology, and drug screening.

Taken together, using the 2D hiPSC-CMs and 3D human cardiac organoids, we have identified the ncRNA IGBP1P1, as a pathologic stress-induced modulator of cardiomyocyte hypertrophy and contractile function. IGBP1P1 depletion rescued cardiomyocyte size and improved cardiac contractility. Thus, blocking the ncRNA IGBP1P1 could be a promising strategy to improve cardiac function in cardiovascular disease.

## Acknowledgments

C.Z., N.B., T.Y., M.S. and J.K. planned and designed the experiments and wrote the paper. N.B., F.B.A. and M.S. performed the bioinformatics analysis. C.Z. executed most of the experiments with help from M.W., Y.W., A.P.D., M.D. and S.D.. J.K., T.T.D. and M.D.P. provided cells and material for 2D and 3D experiments.

## Sources of funding

This work was supported by SFB-TRR 267 (Non-coding RNA in the cardiovascular system) and the German Research Foundation (Exc2026) to J.K., M.S. and S.D. and the European Innovation Council (GA: 822455) to J.K., the LOEWE Center for Cell and Gene Therapy to S.D. and J.K., the European Research Council (Angiolnc) to S.D.. C.Z. and Y.W. were supported by the China Scholarship Council (CSC) Grant #202008080200 and #202108080020.

## Disclosures

The authors have nothing to disclose.

## Figure Legends

**Supplementary Figure 1.**
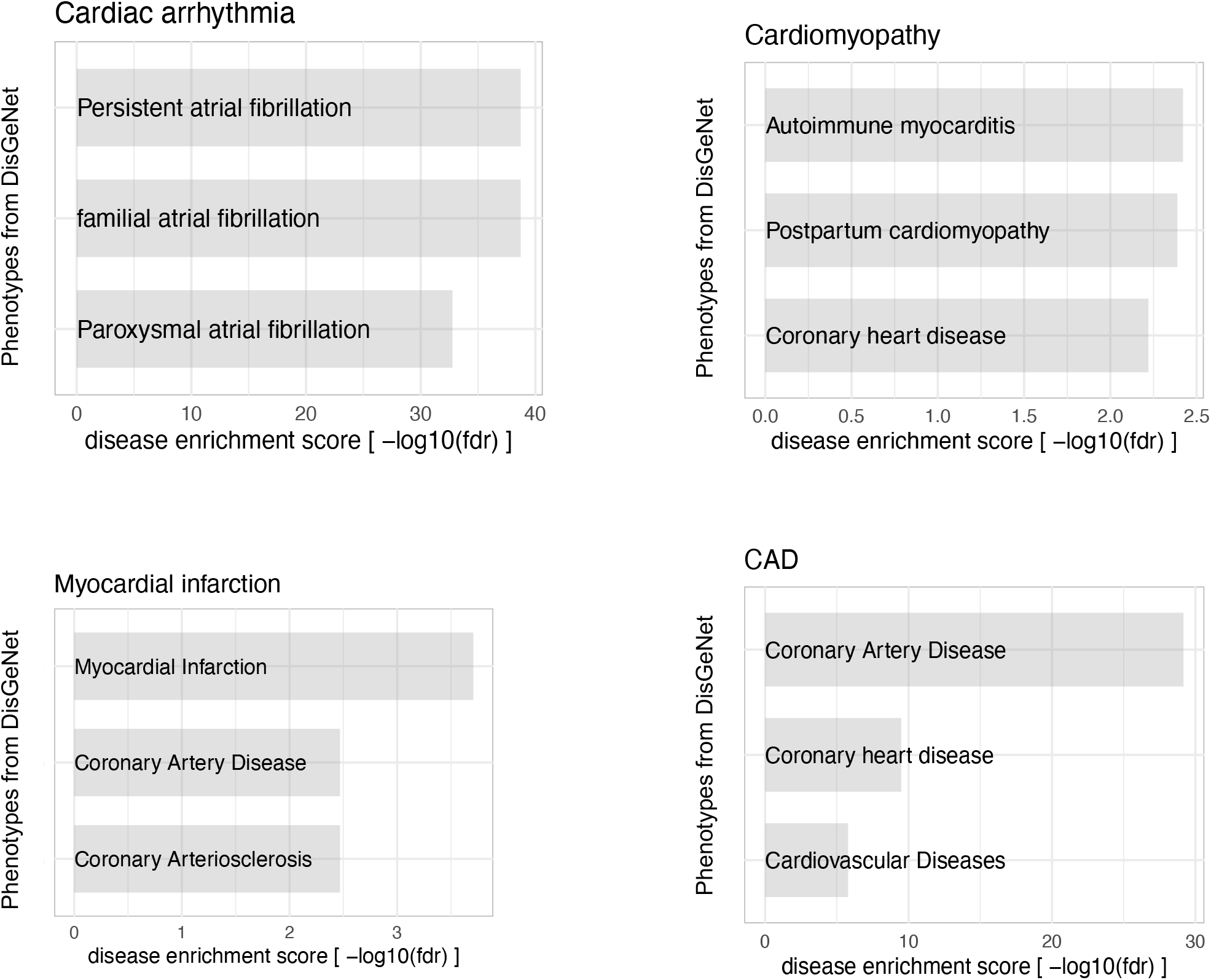
Disease enrichment analysis for protein-coding genes. Barplots representing per GWAS selected phenotypes enriched for protein coding genes identified with SNEEP. The x-axis shows the -log10 FDR corrected p-value of the disease enrichment analysis performed with the DisGeNet software (usage of DisGeNET similar to method section ‘Identification of disease associated genes using rSNPs’, as input the protein-coding genes from the SNEEP result per GWAS are taken) (see also Table S1). For the GWAS Myocardial ischemia and Aortic stenosis the disease enrichment analysis was not possible, because only 5 and 12 protein coding genes were associated.

**Supplementary Figure 2.**
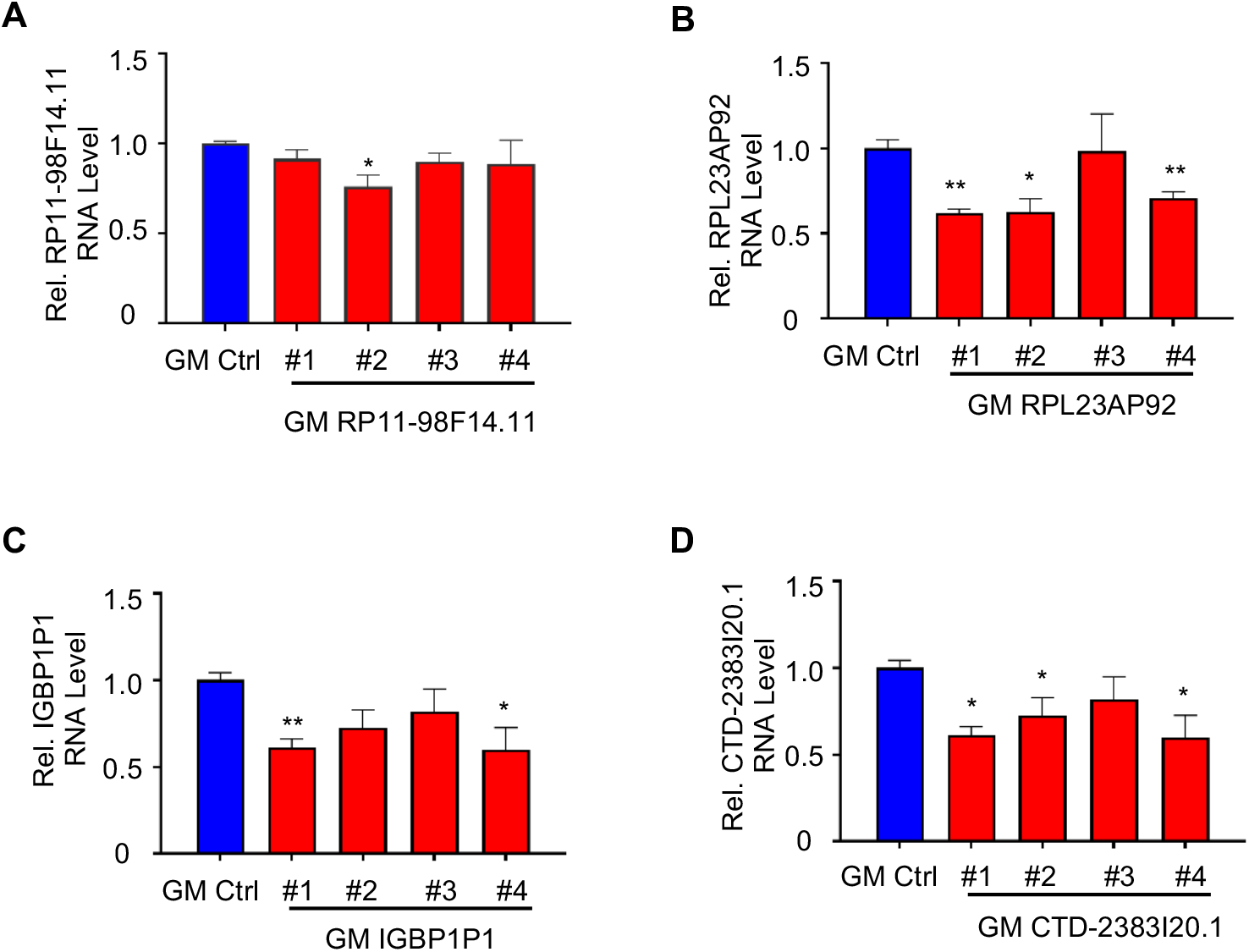
Identification of the best GapmeRs in hiPSC-CMs. (A) Relative RNA expression level of *RP11-98F14*.*11* in hiPSC-CMs treated with GM RP11-98F14.11 #1, #2, #3, #4 or GM GapmeR Ctrl. Data are normalized to hiPSC-CMs expressing GM Ctrl. Data are represented as Mean±SEM; n=3; \**P*< 0.05. (B) Relative RNA expression level of *RPL23AP92* in hiPSC-CMs treated with GM RPL23AP92 #1, #2, #3, #4 or GM Ctrl. Data are represented as Mean±SEM; n=3; \**P*< 0.05. (C) Relative RNA expression level of *IGBP1P1* in hiPSC-CMs treated with GM IGBP1P1 #1, #2, #3, #4 or GM Ctrl. Data are represented as Mean±SEM; n=3; \**P*< 0.05. (D) Relative RNA expression level of *CTD-2383I20*.*1* in hiPSC-CMs treated with GM CTD-2383I20.1 #1, #2, #3, #4 or GM Ctrl. Data are represented as Mean±SEM; n=3; \**P*< 0.05. Two-tailed unpaired t-test.

**Supplementary Figure 3.**
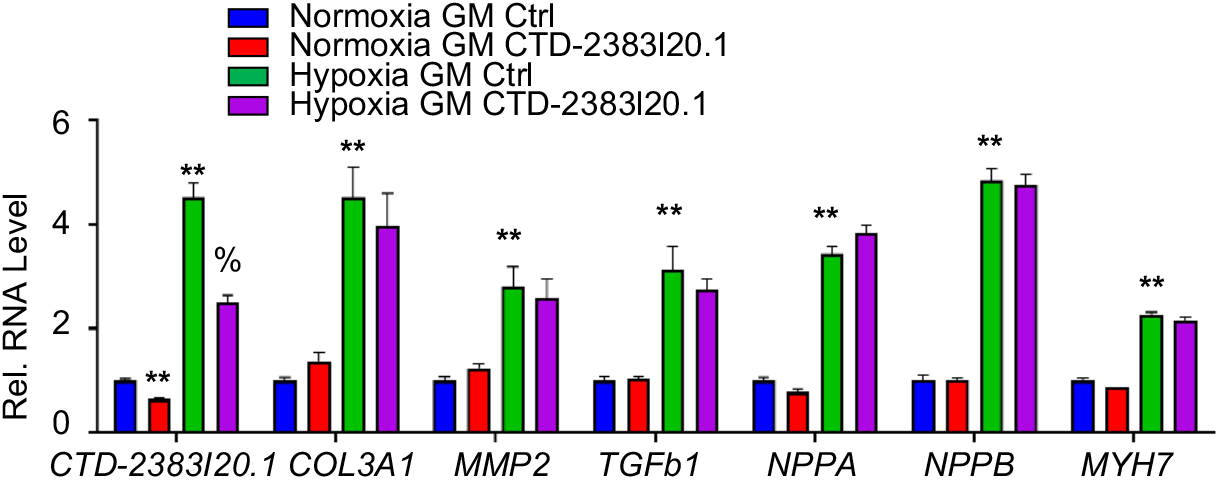
Characterization of CTD-2383I20.1 in hiPSC-CMs. Relative RNA expression of *CTD-2383I20*.*1, COL3A1, MMP2, TGFb1, NPPA, NPPB* and *MYH7* in hiPSC-CMs transduced with either GM CTD-2383I20.1 or GM Ctrl in both normoxia and hypoxia. Data are represented as Mean±SEM; n=3; *, %*P*< 0.05, \*\**P*<0.01.

**Supplementary Figure 4:**
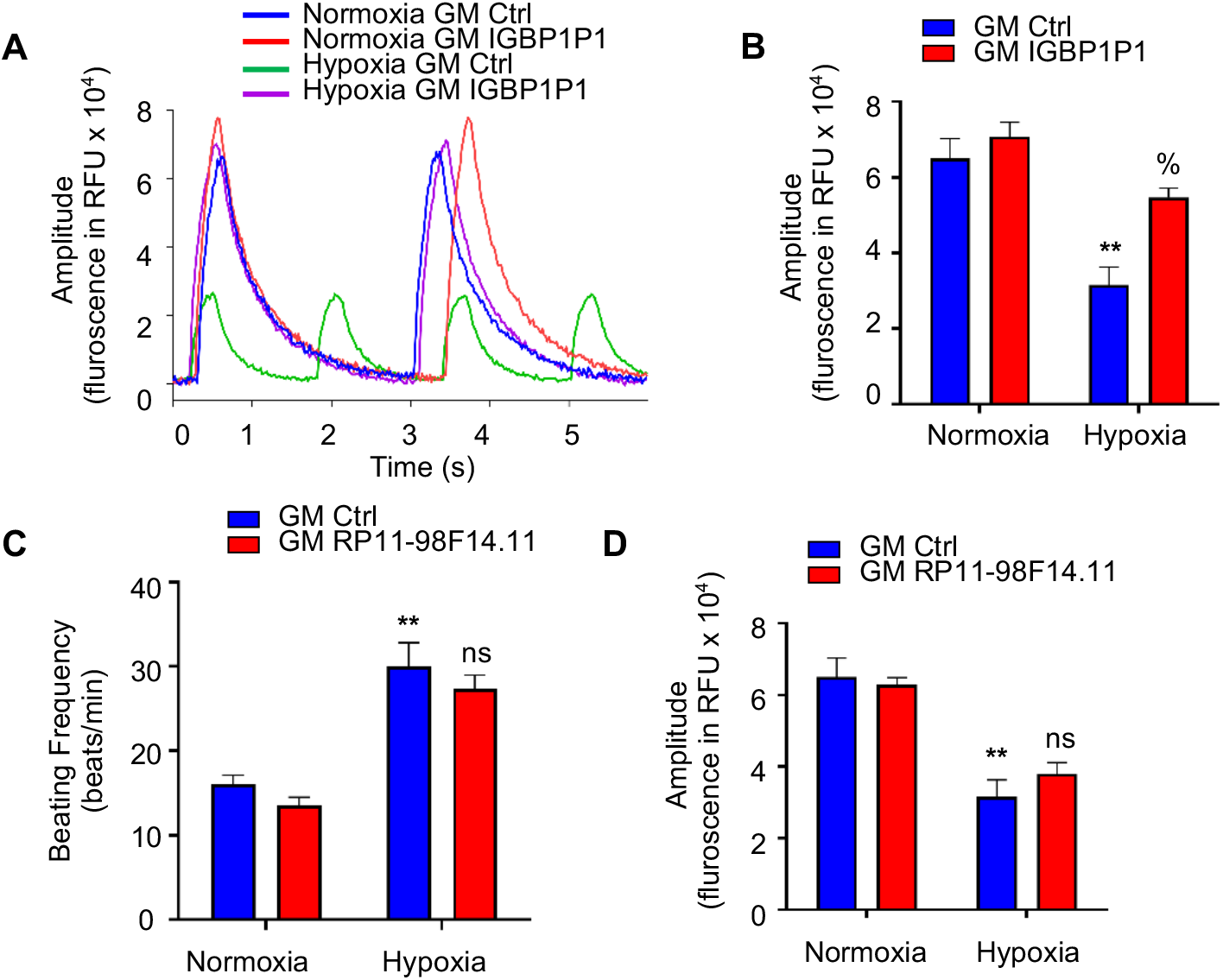
IGBP1P1 inactivation improved contractility in human MCOs. Human MCOs were treated with GM IGBP1P1 or GM Ctrl in both normoxia and hypoxia for 3 days, and then the contractility assays were performed by calcium transient. (A) Representative traces of contractile MCOs. (B) Data are represented as Mean±SEM; *, %*P*< 0.05. Two-tailed unpaired t-test. (C) Human MCOs were treated with GM RP11-98F14.11 *o*r GM Ctrl in both normoxia and hypoxia for 3 days, and then contraction was determined by counting beats per minute. (D) The contractility assays were performed by calcium transient. Data are expressed as means ± SEM. \**P*<0.05, \*\**P*<0.01. Two-tailed unpaired t-test.

## Notes

### Competing Interest Statement

The authors have declared no competing interest.

