## Supplementary tables S8-S10 for "CVD-associated SNPs with regulatory potential drive pathologic non-coding RNA expression"

### Table S8 : List of ncRNAs

| Gene ID | Gene Name | Related Human Disease |
|---|---|---|
| ENSG00000232978 | RP11-146N23.4 | Aortic Stenosis |
| ENSG00000273226 | RP11-513M16.8 | Aortic Stenosis |
| ENSG00000274220 | RP11-77K12.9 | CAD |
| ENSG00000233471 | KRT18P62 | CAD |
| ENSG00000246889 | AP000487.5 | CAD |
| ENSG00000259999 | RP11-252K23.1 | CAD |
| ENSG00000262583 | RP11-77K12.5 / TMEM231P1 | CAD |
| ENSG00000250474 | WBP1LP2 | CAD |
| ENSG00000178803 | ADORA2A-AS1 | CAD |
| ENSG00000201151 | SNORD56 | CAD |
| ENSG00000244691 | RPL10AP1 | CAD |
| ENSG00000248278 | SUMO2P17 | CAD |
| ENSG00000258978 | HIF1AP1 | CAD |
| ENSG00000128262 | POM121L9P | CAD |
| ENSG00000283828 | RP11-98F14.12 | Cardiac Arrythmia |
| ENSG00000238228 | OR7E7P | Cardiac Arrythmia |
| ENSG00000250899 | RP11-253E3.3 | Cardiac Arrythmia |
| ENSG00000271146 | RP11-479F13.1 | Cardiac Arrythmia |
| ENSG00000252072 | RNA5SP320 | Cardiac Arrythmia |
| ENSG00000269125 | RP11-98F14.11 | Cardiac Arrythmia |
| ENSG00000270723 | RPL23AP92 | Cardiac Arrythmia |
| ENSG00000274415 | RP11-433J22.2 | Cardiac Arrythmia |
| ENSG00000272954 | KB-1440D3.13 | Cardiomyopathy |
| ENSG00000226677 | IGBP1P1 | Cardiomyopathy |
| ENSG00000274281 | RP11-326N17.2 | Cardiomyopathy |
| ENSG00000223534 | HLA-DQB1-AS1 | Cardiomyopathy |
| ENSG00000227409 | ZMYM4-AS1 | Cardiomyopathy |
| ENSG00000229391 | HLA-DRB6 | Cardiomyopathy |
| ENSG00000252042 | Y RNA | Cardiomyopathy |
| ENSG00000259345 | RP11-624L4.1 | Cardiomyopathy |
| ENSG00000263786 | RP11-649A18.4 | Cardiomyopathy |
| ENSG00000275293 | Metazoa SRP | Cardiomyopathy |
| ENSG00000249994 | CTD-2383I20.1 | Myocardial Infarction |
| ENSG00000225124 | RPL23AP36 | Myocardial Infarction |
| ENSG00000272750 | RP11-378J18.8 | Myocardial Infarction |
| ENSG00000229116 | RP11-20J15.3 | Myocardial Infarction |
| ENSG00000265215 | MIR4269 | Myocardial Infarction |
| ENSG00000266936 | CTC-215O4.4 | Myocardial Infarction |
| ENSG00000233579 | KRT8P15 | Myocardial Infarction & CAD |
| ENSG00000270001 | RP11-218C14.8 | Myocardial Ischemia |

#### Table S9: List of qRT-PCR primers used in the study

| Target gene name | Forward primer | Reverse primer |
|---|---|---|
| RP11-146N23.4 | TGTTTTGTCTCCGGTGTTCCA | ATAAGGCCAGGTGCGGTG |
| RP11-513M16.8 | TATCTGCGCCTTAACCAGACC | AATGCAAGCTCTTTGTTGGCA |
| RP11-77K12.9 | CCATGAAAGTCGGCCCAAGA | GCGTAGGGTCACACTCTTCC |
| KRT18P62 | GCCTTCATCGTTCTGCACAC | CATGGATGTCGCTCCTCACA |
| AP000487.5 | GAGCCTCTTCATCTCTCCATCC | CACGCATGTGGCCCTTTCT |
| RP11-252K23.1 | GACCAACGTTTCTTGGCTTGA | GCTGGGACTCAGAAGTTGCTT |
| RP11-77K12.5 / TMEM231P1 | AGGCAGAGCCATGAAACTCC | CACCCAGGCGAACTTTACCA |
| WBP1LP2 | TACAGTGACTTCCAGCTACGC | CTTGGGGGTCTTGTGATGCT |
| ADORA2A-AS1 | GCACCACATGCTTGTCTACGA | ATAGAGTCAGGGTTCCAGGCA |
| SNORD56 | ATGTCAATAGTTTTCATCAACAGCA | CCACTCAGACCCAAAGTATCGAC |
| RPL10AP1 | CCCACGAATCCTCGGCATAG | TGCACAAGCTCATCGTCTGT |
| SUMO2P17 | TGAGGTAGATCAGATTCCCATTC | TATCTTCATCCTCCATTTCCAACT |
| HIF1AP1 | TGGAACATTATTAACAGCAGCCAG | TTGCATTCTTTTACACGTTTCTAGG |
| POM121L9P | CCAGCATCTTATTAGAGGACGGAA | TCCCTGAGGACTCTAGCACG |
| RP11-98F14.12 | CCTGGATGCAGGCATGCTAA | GACCTGACCTGGCACAGTTG |
| OR7E7P | ATGGTGTAGTGGCGTCAGTG | CTCCGCAGGGCACTTTGTAT |
| RP11-253E3.3 | CACCTAGTGGCTCTTTGGGG | GAGTGCCAGACACACGGTAA |
| RP11-479F13.1 | TGGACATGGGATTGGTTGAGT | GTTTCTCATGCTGAGAGTGGC |
| RNA5SP320 | TGAACACAAATGCGCAGAGT | AGTTCTCAGTTCATCTCCCATCC |
| RP11-98F14.11 | GGAGGTCCTGTAGATCCGGT | GCTGAGAAGGCGCTGATTTT |
| RPL23AP92 | GCGACCAACAAGTTCTCCCA | GCCCTGATCACGGTGTTGAT |
| RP11-433J22.2-transcript201 | AACTGTAAAGGAGCTGCAGGG | GCCCTGGGGGAAAATTCTTGG |
| RP11-433J22.2-transcript202 | GGCCATCTCACCACTACTCC | CACAGACAACCTGATCACCCT |
| KB-1440D3.13 | ATCTCTTGTGCCCACCTTGAG | ACCTCCTTAGTTCCCAGCGT |
| IGBP1P1 | GATCAGGGAATAGCCAAGGCA | CTCTGTTGGCTGCCATAGTC |
| RP11-326N17.2 | AGTGCTGGTTACCAACTTTCCT | GGAGTGCCAAGATCGCATGA |
| HLA-DQB1-AS1 | CAGCTTGATGCAGATGTGTGG | CATGATGGTGGCTACTGCCT |
| ZMYM4-AS1 | GACATGCTGTCAAGGGTAGGA | CCAGACTGACCTTATCATTGTGGT |
| HLA-DRB6 | TTGGAGCAGGCTAAGTGTGAG | TCCGTAACTGCCTGGAACTC |
| Y RNA | GCTGGTAGTGAGTTATCTTG | ACAGACTAGCCAAGTGCAGTA |
| RP11-624L4.1-transcript 1-4 | ACCAGAAGCACTCCAAGAACAA | GACTTTGAAGTGACAGGCTGG |
| RP11-624L4.1-transcript 5 | ATGCAGCCATCAGCCTCAAT | GCTTTTGGGTTGGTTGCCTT |
| RP11-624L4.1-transcript 6,9 | CCTGCTGTGGGAGTAACCAT | AAGCCCTAGAGGGACAAGGT |
| RP11-624L4.1-transcript 7-8 | TGGCCTGCATCCACTGTCT | AGACCCAAGATGGCCGAATAGG |
| RP11-649A18.4 | TCAAGCTCAGCTCACAGCAT | TCACCAAGCAGGTAACCAATGT |
| Metazoa SRP | ATACTGATGGGGTGTCTGCA | GTCCCGAACTCCTGACCTC |
| CTD-2383I20.1 | GACTCTGGCCTGAAGAAAGCA | TTGGCTCTCGGTGATCCTACT |
| RPL23AP36 | GCTAAAAGGCATCCACCCCA | GGCTGCGTTTGGAACCATAG |
| RP11-378J18.8 | AATGACCGCTCTGTCTTCTGT | AGGGTACAGTTGTTAGGGTAACG |
| RP11-20J15.3 | CCTGTGACCCGGATCCAAC | CAAATGAACAGAAGCTGGGGG |
| MIR4269 | CCTGCAGGCACAGACAGC | CCATCCCAGGCCTGACAGA |
| CTC-215O4.4 | AGGCCGCATTAAGAGCATGA | GTAGCAGGTCCTGTGTGAGG |
| KRT8P15 | TTCGGCAACTGCTCCTATGC | CACTGGCCCCACCATAACTT |
| RP11-218C14.8 | GGGCCTGTAAATGCCTCCC | AGATGAAAGGTGCAAGGGCG |
| COL3A1 | TTGAAGGAGGATGTTCCCATCT | ACAGACACATATTTGGCATGGTT |
| MMP2 | TACAGGATCATTGGCTACACACC | GGTCACATCGCTCCAGACT |
| TGFb1 | CAATTCCTGGCGATACCTCAG | GCACAACTCCGGTGACATCAA |
| NPPA | CAACGCAGACCTGATGGATTT | AGCCCCCGCTTCTTCATTC |
| NPPB | TGGAAACGTCCGGGTTACAG | CTGATCCGGTCCATCTTCCT |
| MYH7 | GACCAGATGAATGAGCACCG | GGTGAGGTCGTTGACAGAAC |
| MYH6 | CTCCTCCTACGCAACTGCCG | CGACACCGTCTGGAAGGATGA |
| RP11-98F14.11 | TAAATTGAAGCACGCGGAGAG | GCTGACCTCTGAGAAGCGTG |
| RPL23AP92 | GCGACCAACAAGTTCTCCCA | GCCCTGATCACGGTGTTGAT |
| IGBP1P1 | GATCAGGGAATAGCCAAGGCA | CTCTGTTGGCTGCCATAGTC |
| CTD-2383I20.1 | TGGCCTCATCAGAACCAAGAC | CAAGCTGGCCATTTTGTTTGT |
| HPRT | CCTGGCGTCGTGATTAGTGAT | AGACGTTCAGTCCTGTCCATAA |

### Table S10: Characteristics of the LNA GapmeRs used in the study

| ID | Sequence (5'-3') |
|---|---|
| GapmeR negative Control | AACACGTCTATACGC |
| GapmeR RP11-98F14.11 #1 | CGCGTGCTTCAATTTA |
| GapmeR RP11-98F14.11 #2 | CTCCGCGTGCTTCAAT |
| GapmeR RP11-98F14.11 #3 | CTGAGAAGGCGCTGAT |
| GapmeR RP11-98F14.11 #4 | TCGCTGAGAAGGCGCT |
| GapmeR RPL23AP92 #1 | TTGGTCGCATAGTGGT |
| GapmeR RPL23AP92 #2 | TTTTGACCTCGACATC |
| GapmeR RPL23AP92 #3 | CGACATCCACAATGAA |
| GapmeR RPL23AP92 #4 | GCTTTTTGACCTCGAC |
| GapmeR IGBP1P1 #1 | TGGCGAGATGAATTAG |
| GapmeR IGBP1P1 #2 | GGATAAGCCATGGAGA |
| GapmeR IGBP1P1 #3 | GAAATTAGCAGTGTGA |
| GapmeR IGBP1P1 #4 | GCAATGAGGCTAGGAT |
| GapmeR CTD-2383I20.1 #1 | GAAGGTTGGTCGTCTT |
| GapmeR CTD-2383I20.1 #2 | GTTGGTCGTCTTGGTT |
| GapmeR CTD-2383I20.1 #3 | CTTGCCAGCATCTGAT |
| GapmeR CTD-2383I20.1 #4 | AACAAGCTGGCCATTT |
